# A characterization of the electrophysiological, morphological and input domains of vasoactive intestinal peptide (VIP) interneurons in the medial entorhinal cortex (MEC)

**DOI:** 10.1101/2020.05.15.097972

**Authors:** Saishree Badrinarayanan, Frédéric Manseau, Byung Kook Lim, Sylvain Williams, Mark P. Brandon

**Affiliations:** Department of Psychiatry, Douglas Hospital Research Centre, McGill University, Montreal, QC, Canada; Integrated Program in Neuroscience, McGill University, Montreal, QC, Canada; Department of Biological Sciences, University of California San Diego, San Diego, CA, USA

## Abstract

Circuit interactions within the medial entorhinal cortex (MEC) translate movement into a coherent code for spatial location. Entorhinal principal cells are subject to strong lateral inhibition, suggesting that a disinhibitory mechanism may drive their activation. Cortical Vasoactive Intestinal Peptide (VIP) expressing inhibitory neurons predominantly contact interneurons, providing a local disinhibitory mechanism. Here, we investigate the electrophysiological and morphological properties of VIP cells using *in vitro* whole-cell patch clamp recordings and use rabies-mediated circuit tracing to discover long-range inputs that may modulate this population in mice. We report physiological and morphological properties of VIP cells that differ across lamina and along the dorsal-ventral MEC axis. Furthermore, we reveal long-range inputs to VIP neurons from regions known to encode proprioceptive and auditory information, including the mesencephalic trigeminal nucleus and superior para-olivary nuclei, respectively. These results characterize the properties of VIP cells and reveal sensory modalities that could drive disinhibition in the MEC.

## Introduction

The medial entorhinal cortex (MEC) is a 6-layered cortical structure involved in episodic memory and spatial navigation. Layers 1-3 (LI-III) form the input and layers 5-6 (LV-VI) form the output domain to various cortical structures (Ramsden et al. 2015; Sürmeli et al. 2015; Witter et al. 2017) and for the hippocampus, projections from the MEC originate from the superficial layers and outputs from the hippocampus terminate in the deep layers (Canto, Wouterlood, & Witter, 2008). Through these reciprocal connections, these regions work together to support circuits responsible for spatial navigation and memory (Moser, Kropff, & Moser, 2008). The MEC consists of several spatially tuned cells (Hafting et al. 2005; Solstad et al. 2008; Sargolini et al. 2006) of which, ‘grid cells’ and ‘border cells’ are hypothesized to emerge locally. Grid cells fire in a distinct hexagonal pattern that span the entire environment an animal traverses. Navigational information like an animal’s displacement, trajectory, or angular motion can be decoded from grid cell populations (Moser et al., 2014). Using this information, as pointed by numerous theoretical and computational studies, grid cells are suggested to perform a function known as path integration (Mittelstaedt and Mittelstaedt 1980; McNaughton et al. 2006). Border cells fire reliably when the animal is in close proximity to one or more edges of an environment (Solstad et al., 2008), providing information of environment boundaries. Thus, the MEC is hypothesized to track movement through space, as well as the spatial extent of its environment.

Several lines of evidence indicate that grid cells and border cells are under strong lateral inhibition. While both cell types are observed across MEC lamina, grid cells are particularly abundant in layer II of the MEC (Sargolini et al. 2006, Boccara et al. 2010). *In vitro* experiments using paired and multiple whole-cell recording techniques, have shown that excitatory coupling in LII is present but sparse, and instead, the excitatory cells are connected via strong lateral inhibition (Couey et al., 2013; Fuchs et al., 2016; Winterer et al., 2017). Consistent with this, *in vivo* optogenetic activation of parvalbumin (PV) expressing inhibitory neurons directly inhibits both grid cells and border cells, with grid cells providing a back projection to PV cells (Buetfering, Allen, & Monyer, 2014). Finally, DREADD inactivation of the PV inhibitory cell populations, but not the somatostatin inhibitory populations (SOM), disrupt the spatial firing pattern of grid cells, while manipulations to both these cell populations does not affect border cells (Miao, Cao, Moser, & Moser, 2017). Computational attractor models have shown how excitation-inhibition-excitation (E-I-E) networks or recurrent inhibitory networks can generate grid cells (Bonnevie et al. 2013, Schmidt-Hieber and Nolan 2017; Kang and Balasubramanian 2019). The role of excitatory and certain inhibitory cell populations has been studied extensively, but the interplay required to generate and maintain this Excitation/Inhibition (E/I) balance remains elusive.

While studies have shown that the interneurons within the MEC receive inhibitory projections from the medial septum (Fuchs et al., 2016; Gonzalez-Sulser et al., 2014) another form of disinhibition conserved across cortical structures, is mediated by the Vasoactive Intestinal Peptide (VIP) expressing interneurons. These cells target both PV and SOM inhibitory cells (Kepecs and Fishell 2014). Pyramidal cells have also been noted to be a minor monosynaptic target of VIP interneurons (Batista-Brito et al. 2017; Rhomberg, Rovira-Esteban, et al. 2018; Krabbe et al. 2019). Thus, in terms of cortical connectivity, VIP interneurons can provide for transient and selective breaks in the E/I balance by disinhibiting excitatory cells in other cortical regions (Letzkus, Wolff, & Lüthi, 2015). The disinhibitory circuit motif has been well defined in the hippocampus (Tyan et al. 2014; Turi et al. 2019), auditory cortex (Pi et al., 2013), basolateral amygdala (Krabbe et al., 2019; Rhomberg et al., 2018), visual cortex (Batista-Brito et al., 2017) and somatosensory cortex (Prönneke et al., 2015). In addition to this, these studies show VIP interneurons exhibit a range of molecular, electrophysiological and morphological characteristics, all of which remain ill-defined in the MEC.

Here, we provided a detailed account of the anatomical and *in vitro* physiological properties of VIP cells in the MEC. We crossed VIP-ires-cre with tdTomato Ai9 mice to analyze the various properties of VIP interneurons. We performed immunohistochemistry (IHC) against several molecular markers to validate and elucidate the molecular composition of VIP interneurons in the MEC. We targeted individual VIP neurons in all layers and along the dorsal-ventral (D-V) axis of the MEC using whole-cell patch-clamp recordings to ascertain their electrophysiological and morphological profiles and compared them with other brain structures. To investigate the areas and cell types that can activate VIP cells, we visualized brain-wide inputs to these cells in the MEC using retrograde rabies virus tracing. Our findings provide for an overview of the anatomical characteristics of these interneurons within the MEC.

## Results

### Molecular characterization of VIP cells in the MEC

We quantified the distribution of MEC VIP neurons in layer-specific and layer-independent manners by crossing transgenic VIP:ires:cre and tdTomato cre-reporter mice (n = 3). A total of 6219.56 ± 607.81 cells expressed tdTomato (tdTom), of which 5849.51 ± 633 cells were labelled with VIP IHC (Fig. 1C). Thus, 94.05% of tdTom cells colocalized with VIP (VIPtdTom). When quantified by cortical layer, VIP cell bodies were not equally distributed. At 89.28%, superficial MEC (LI-III) had the highest proportion of VIPtdTom cell bodies, while the deep MEC (LIV-VI) consisted of 19.71% of cell bodies (4673 ± 388.40 vs 1153.43 ± 251.21) (Fig. 1D). These results indicate that tdTom expression is highly specific (Fig. 1A-D) for VIP-positive cells in the VIPcre/tdTom mouse line and thus suitable for the study. To ascertain if VIP cells express other molecular markers, we performed IHC against antibodies for GABAergic cells (Fig. 1E-E”), Calretinin (CR, Fig. 1H-H”) and estimated the percent of colocalization with tdTom.

**Fig 1.**
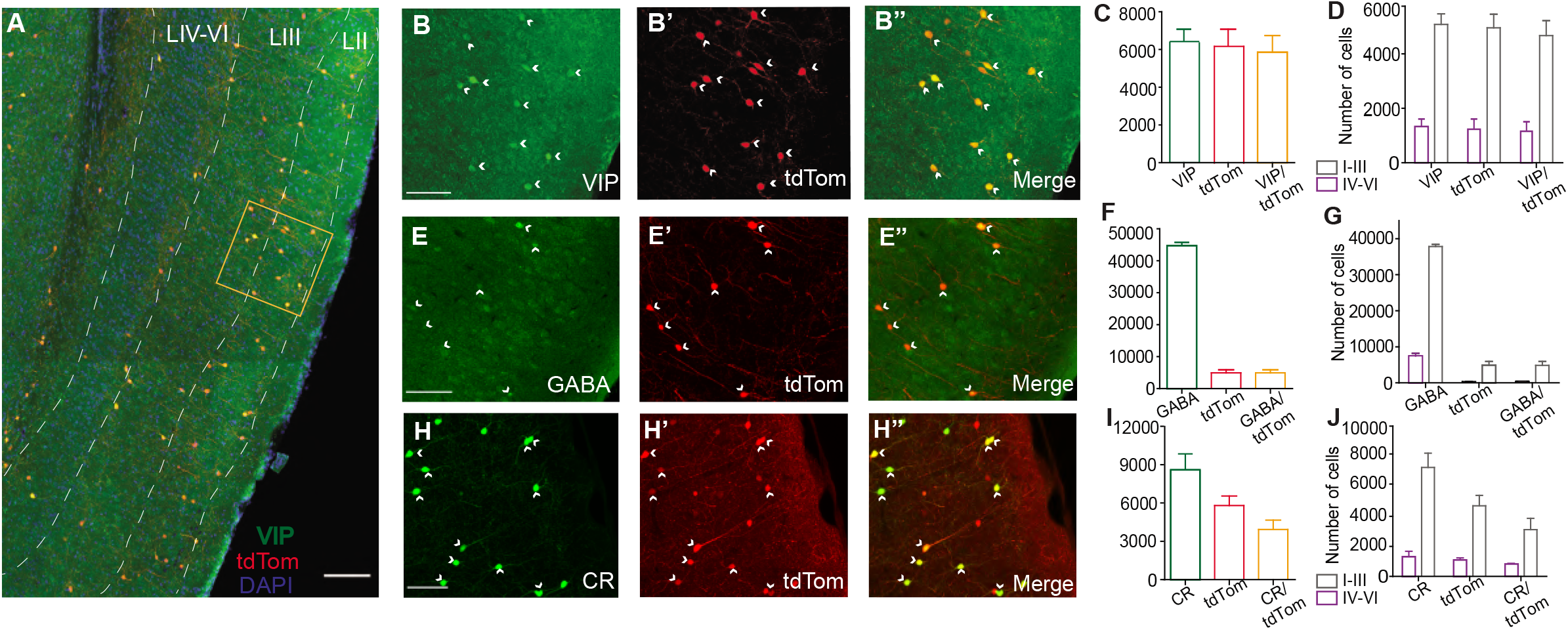
Specificity of cre-driven fluorescence for VIP in VIPcre/tdTom mice and presence of colocalization with other molecular markers. (A) Co-expression of VIP (green) and tdtom (red) in a sagittal section of the MEC at low maginification 10X. Layers of the MEC (white dashed lines) marked using DAPI as reference. High maginification of framed area (orange box) is shown in B-B”. (B-K”) 20X maximum projection intensities of a 40-μm thick section of superficial MEC treated with IHC using αVIP AB (B-B”), αGABA AB (E-E”) and αCR AB (H-H”). Fluorescence of cre-driven tdTomato expression is shown in red (B’-H’) and colocalization of the two channels is shown in (B”-H”). Presence of colocalization is indicated by yellow cell bodies and white arrow heads. Graphs depicting the total number of cells present in the MEC (C, F and I) and distribution of somas in different layers of the MEC (D, G and J) for αVIP, αGABA and αCR respectively are shown here. Each bar represents a mean and error bars represent SEM. For panels B-H” scale bars are 250μm. Scale bar in A is 500μm.

We observed that all tdTom cells colocalized with GABA (4945.75 ± 685.01 cells). Our results indicate that 11.05% of the total GABAergic population are VIP+ (4945.75 ± 685.01 of 44726.64 ± 727.24 cells) (Fig. 1F), similar previous reports in the cortex (Rudy 2013; Tremblay, Lee, and Rudy 2016; Prönneke et al. 2015). Much like other structures, we also found VIP cells to express Calretinin (CR). Of all the CR-expressing cells, 45.63% colocalized with tdTom (3922 ± 521.05 of 8594.54 ± 682 cells) while 67.59% of tdTom cells colocalized with CR (3922 ± 521.05 of 5802 ± 520.72 cells; CRtdTom) (Fig. 1I). This is in line with results of VIP cells in hippocampus(Tyan et al., 2014) and basolateral amygdala (Rhomberg et al., 2018), but much higher than the 35% of colocalization observed in the visual cortex (Gonchar, Wang, & Burkhalter, 2008). We further looked at the distribution of CRtdTom cells across the layers (Fig. 1J). A high density of CRtdTom cells (79.27%) were found in the superficial layers (3109.44 ± 534.83 cells). Previous studies reported that VIP+ cells do not overlap with other molecularly distinct subpopulations such as PV and SOM (Prönneke et al. 2015). We observed the same in our experiments (Fig S1. A-B”).

### Electrophysiological properties of VIP neurons vary between superficial and deep layers of the MEC

Next, we determined the electrophysiological properties of 51 VIP cells from all layers of the MEC (LI-III n = 29; LIV-VI n = 22). Two passive membrane properties were significantly different between the two populations (Fig. 2B): VIP neurons in LIV-VI had higher input resistance (Fig. 2B(i)) compared to neurons present in LI-III (528.68 ± 25.41 vs 432 ± 20.51 MΩ, Mann Whitney U = 172.0, p = 0.0053). Consistent with this difference, the amplitude in threshold current (rheobase) (Fig. 2B(ii)) required to generate an action potential in deep MEC was significantly less than the superficial VIP cells (20.79 ± 2.85 vs 39.08 ± 4.24 pA, Mann Whitney U = 161, p = 0.0024). Layer differences were also observed for active membrane properties (Fig. 2C): Action potential from cells in the superficial layers had a smaller half-width compared to VIP cells in the deep layers (1.24 ± 0.098 vs 1.52 ± 0.103 ms, Mann Whitney U = 199, p = 0.0230) (Fig. 2C(i)), accordingly significant difference was observed in the maximum rise slope of spikes (Fig. 2C(ii)) between superficial and deep VIP cells (175.12 ± 12.67 vs 131.26 ± 8.81 mV/ms, Mann Whitney U = 201, p = 0.025). We also found deep VIP cells to have a much longer duration in AHP time compared to superficial cells (0.079 ± 0.0042 vs 0.064 ± 0.010 s, Mann Whitney U = 154, p = 0.0018) (Fig. 2C(iii)). No significant differences were observed in properties such as resting membrane potential (population average of −60.11 ± 1.28 mV), voltage sag amplitude (3.85 ± 0.33 mV), firing threshold (−40.32 ± 0.80 mV), peak amplitude (64.58 ± 1.34 mV), half-amplitude (32.29 ± 0.67 mV), AHP amplitude (−11.10 ± 0.76 mV), peak inter-spike interval (51.15 ± 4.83 Hz) and steady state inter-spike interval (33.78 ± 3.71 Hz) (Table 2).

**Fig 2.**
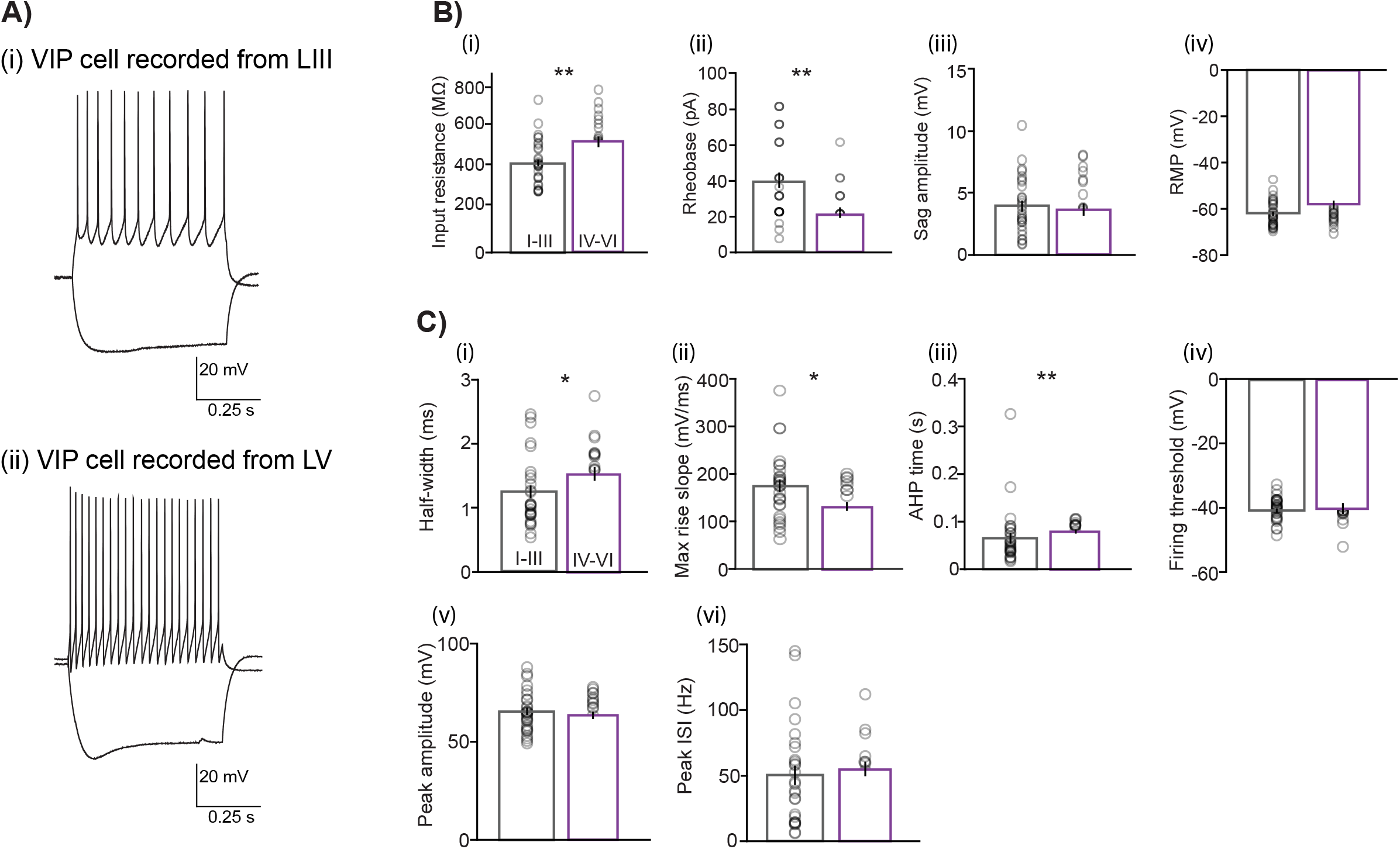
Comparison of electrophysiological properties between LI-LIII and LIV-LVI VIP cells. (A) Examples of membrane potential responses to current steps (−80pA for lower trace and +80pA for upper trace) recorded from VIP cells located in the (i) superficial (LIII) and (ii) deep (LV) layers of the MEC. (B) Passive and C) Active membrane properties for 51 VIP cells separated into layer I-III (n=29) and layers IV-VI (n=22). Except (iii) sag amplitude and (iv) peak ISI where layer I-III (n=27) and layers IV-VI (n=21). Each bar represents a mean and error bars represent SEM. Significant differences were found in B (i) input resistance, (ii) rheobase and C(i) half-width, (ii) AHP time and (iii) max rise slope. (*= p<0.05, **= p <0.01)

**Table 1.** Tabulation of all passive and active membrane properties measured for VIP cells located in superficial (LI-LIII) and deep (LIV-LVI) layers in the MEC. p values obtained from t-test.

| Properties measured | LI-III (n = 29) | LIV-VI (n = 22) | t test |
| --- | --- | --- | --- |
| Input resistance (MΩ) | 432 ± 20.51 | 528.68 ± 25.41 | p = 0.005 |
| Rheobase (pA) | 39.08 ± 4.24 | 20.79 ± 2.88 | p = 0.002 |
| Sag amplitude (mV) | 4.011 ± 0.46 (n = 27) | 3.63 ± 0.49 (n = 21) | p = 0.692 |
| Decay tau (ms) | 8.66 ± 2.64 | 14.60 ± 4.43 | p = 0.137 |
| RMP (mV) | -61.86 ± 1.06 | -57.80 ± 2.53 | p = 0.212 |
| Half-width (ms) | 1.24 ± 0.09 | 1.52 ± 0.103 | p = 0.023 |
| AHP time (s) | 0.064 ± 0.010 | 0.079 ± 0.004 | p = 0.001 |
| Max rise slope (mV/ms) | 175.12 ± 12.6 | 131.26 ± 8.81 | p = 0.025 |
| AHP amplitude (mV) | -11.04 ± 1.15 | -11.18 ± 0.92 | p = 0.871 |
| Peak amplitude (mV) | 65.47 ± 1.86 | 63.40 ± 1.87 | p = 0.621 |
| Half-amplitude (mV) | 32.73 ± 0.93 | 31.70 ± 0.93 | p = 0.621 |
| Peak ISI (Hz) | 52.83 ± 8.12 (n = 27) | 42.96 ± 6.16 (n = 21) | p = 0.766 |
| Steady state ISI (Hz) | 34.12 ± 5.29 (n = 21) | 25.93 ± 5.64 (n = 18) | p = 0.169 |
| Spike frequency adaptation % | 10.98 ± 19.8 (n = 21) | 33.83 ± 7.01 (n = 18) | p = 0.638 |

**Table 2.** Tabulation of all passive and active membrane properties measured for all VIP cells located across all layers.

| Properties measured | Population average (n = 51) |
| --- | --- |
| Input resistance (MΩ) | 473.71 ± 17.35 |
| Rheobase (pA) | 31.19 ± 2.99 |
| Sag amplitude (mV) | 3.85 ± 0.33 (n = 48) |
| Decay tau (ms) | 11.23 ± 2.46 |
| RMP (mV) | -60.11 ± 1.28 |
| Half-width (ms) | 1.36 ± 0.07 |
| AHP time (s) | 0.07 ± 0.006 |
| Max rise slope (mV/ms) | 156.20 ± 8.70 |
| AHP amplitude (mV) | -11.10 ± 0.76 |
| Peak amplitude (mV) | 64.58 ± 1.34 |
| Half-amplitude (mV) | 32.29 ± 0.67 |
| Peak ISI (Hz) | 51.15 ± 4.83 (n = 48) |
| Steady state ISI (Hz) | 33.78 ± 3.71 (n = 39) |
| Spike frequency adaptation<br>% | 4.08 ± 16.9 (n = 39) |

### Resting and excitable properties of VIP cells vary along the Dorsal-Ventral (D-V) axis of the MEC

We asked if VIP cells show a gradient in their properties along the D-V axis of the MEC (Fig. 3B-D). We recorded 49 cells (LI-III = 27; LIV-VI = 22 cells). and excluded 2 cells due to an inability to quantify the position of the recording pipette. Since we observed layer differences in the electrophysiological properties, we asked if passive and active properties varied as a function of cell’s location in a layer and along the D-V axis. Input resistance increased for deep VIP cells (Fig. 3B(ii)) along the D-V axis (slope = 0.15 ± 0.056 MΩ R^2^ = 0.268, p = 0.0134). Consistent with the lower input resistance in these cells, the rheobase (Fig. 3C(ii)) required to evoke action potentials was much larger for neurons from dorsal locations as compared to that for ventral locations (slope = −0.016 ± 0.006 pA R^2^ = 0.342, p = 0.0181). Interestingly, the firing threshold of these cells showed no dependence on the location of the cell along the D-V axis (Fig. 3D(i)). For cells located in the superficial layers, the firing threshold (Fig. 3D(ii)) steeply increased with cell’s distance from the dorsal border (slope = 0.0039 ± 0.0018 mV R^2^ = 0.249, p = 0.0013), while other properties showed no dependency on the location of the cell along the dorsal border (data not shown).

**Fig 3.**
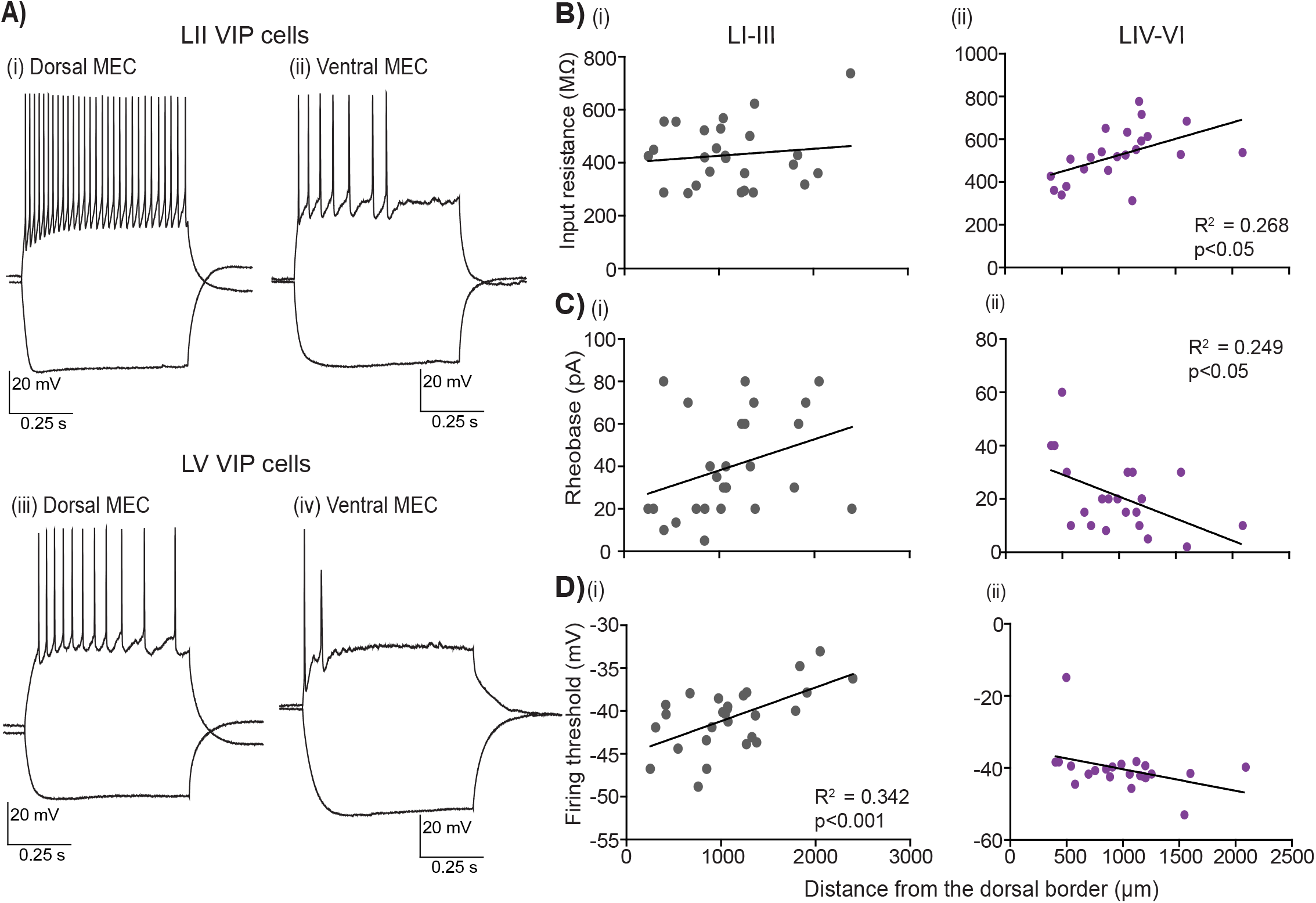
Passive and Active properties as a function of layer and location along the Dorsal-Ventral axis. (A) Examples of membrane potential responses to −80 pA (lower traces) and +80pA (upper traces) current step recorded from LII VIP cells along the D-V axis: (i)dorsal (418.3 μm) and (ii)ventral (1834.4 μm) from the dorsal border. Similarly, example traces from two LV cells recorded from (iii)dorsal (470.2 μm) and (iv)ventral (1546 μm) (B-D) Passive and active properties plotted as a function of location and layer for individual VIP cells recorded from the superficial layers LI-III: 27 cells (B, C and D (i)) and deep layers LIV-VI: 22 cells (B, C, and D (ii)). In all panels, black lines indicate linear fits to the data. For cells located in the deep MEC significance in slope for B (ii)input resistance and C(ii)rheobase is observed, while for cells located in the superficial layers significance in slope for D (i)firing threshold is observed.

### VIP cells show heterogeneity in their firing patterns

As reported in previous studies, VIP cells were never fast-spiking, but showed the following firing patterns, according to description in the Petila convention (Ascoli, 2008): irregular spiking (IS,65%), continuous non-adapting (CNA, 24%) and continuous adapting (CA, 12%) (Fig. S1C). We further investigated the distribution of these firing patterns within the MEC layers. In the superficial layers (n = 29 cells), 76% of the cells had IS firing pattern, while CNA and CA comprised of the remaining 10% and 14% respectively (Fig. 4C). In the deep layers (n = 22 cells), 50% of the cells had IS firing pattern while 41% were CNA and the remaining 9% were CA (Fig. 4D).

**Fig 4.**
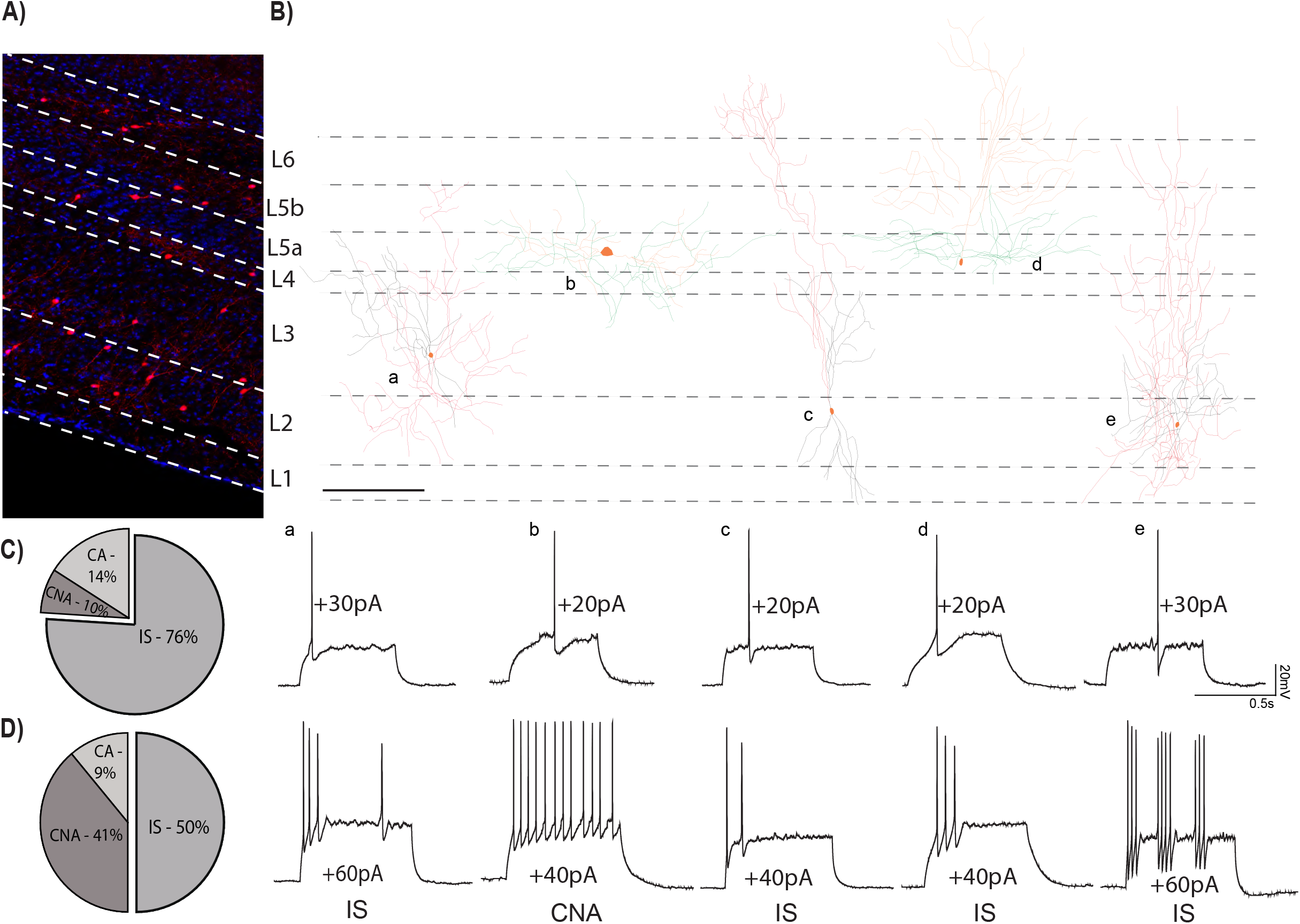
Variability in firing patterns of VIP cells from layers I-III and IV-VI. (A) Distribution of VIP cells (tdTom) in the MEC, the layers have been marked using DAPI as reference. (B) The reconstructed morphology of 5 individual VIP cells is shown in a schematic of the MEC. For all cells soma are coloured in orange. For cells recorded in superficial MEC axonal and dendritic processes are coloured in red and black respectively. For cells recorded from layers IV-VI, axons are in orange while dendrites are coloured green. Below, electrophysiological recordings from the respective cells at rheobase (upper trace) and action potential firing pattern at suprathreshold current (lower traces). Scale bar in B is 100 μm. Percentages of various firing pattern observed in the (C) superficial layers of the MEC and (D) deep layers of the MEC. Abbreviations name the firing pattern after the Petila convention (Ascoli et al.2008; IS, Irregular spiking; CA, Continuous adapting; CNA, continuous non-adapting)

### Morphological features of individual VIP neurons

We next analyzed the dendritic and axonal morphology of 14 VIP neurons from all layers of the MEC were analyzed. In our dataset, we found that most VIP cells presented with a bi-polar/bi-tufted (Fig. 5A(i&ii)) or a multipolar morphology (Fig. 5A(iii)). Our reconstructions indicate that cells within the deep layers showed much denser dendritic arborization within their compartment, compared to the superficial VIP cells. Indeed, we found that cells in the deep layers have a greater horizontal spread of their dendritic trees (Fig. 5C(i)) (385 ± 49.6 vs 205.17 ± 21.43μm, Mann Whitney U = 6.00, p = 0.02), while the dendrites of cells in LI-III are spread across the vertical column (Fig. 5C(ii)) (328.77 ± 19.7 vs 107.66 ± 31.15μm, Mann Whitney U = 1.00, p = 0.0013). We found no significant differences in the horizontal or vertical extent of the axonal arbours between the two groups (data not shown). Given the difference in the dendritic expanse, we asked if the location of the cell influenced its projection to different layers. We saw that cells in the superficial layers extended their axons well into the deep layers, but deep VIP cells made almost no contact to the superficial layers (Fig. 5D(i)). Layer 3 had extensive dendritic arborizations from superficial cells (p<0.001), while dendrites from the deep layers rarely extended beyond L4 and thus their projections were more local (p<0.01) (Fig. 5D(ii)). Sholl analysis revealed that most dendrites and axons were found locally near the soma (≤ 300 μm) for both groups (Fig. 5F).

**Fig 5.**
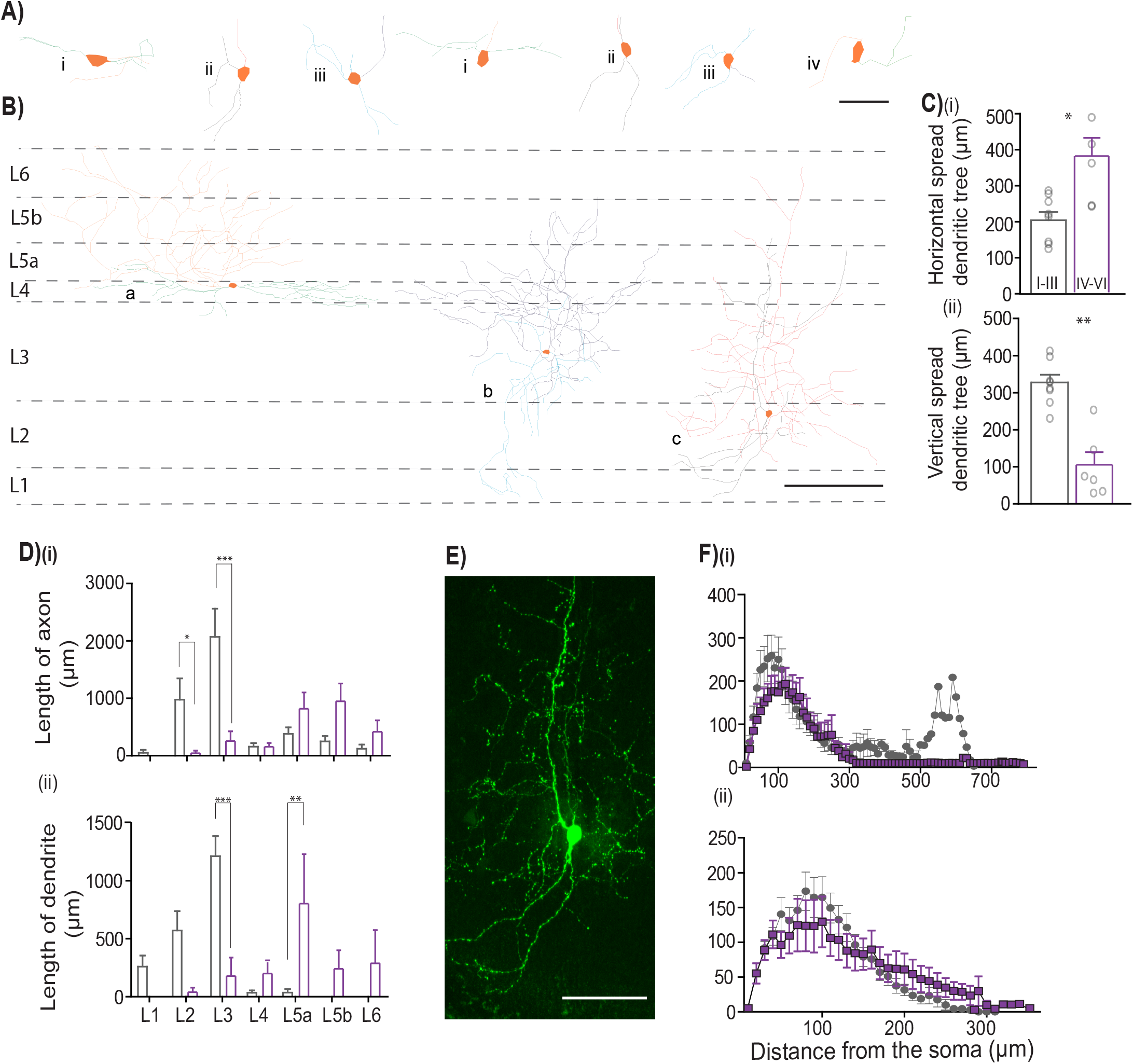
Morphometric analysis of VIP cells. (A) Magnified views of the somatodendritic and axonal processes representing the following morphologies; i – horizontal bi-polar, ii – bi-tufted, iii – multipolar and iv – unipolar architectures. Scale bar are 50μm (B) Schematic of the MEC with neurolucida reconstructions depicting the three predominant morphologies observed in the data; horizontal bi-polar in L4 (a; axons in orange, dendrites in green), multipolar VIP neuron in L3 (b; axons in dark blue and dendrites in light blue) and vertical bi-polar morphology seen in L2 (c; axons in red, dendrites in blue). Scale bars are 100μm. For all cells the soma is coloured in orange. (C) Plots depicting siginificant different features in the vertical (i) and horizontal (ii) spread of the dendritic process Each bar represents a mean and SEM (D) Graphs assessing the total length of process contained in a specific layer for (i) axons and (ii) dendrites Each bar represents a mean and SEM (E) Maximum projection intensity of 207 z-stacks of 0.5 μm depth taken in 20X magnification of a bi-tufted neuron recorded in LII of the MEC filled with neurobiotin (GFP). Scale bar is 250μm (F) Morphological features of dendritic (i) and axonal (ii) arbors obtained from sholl analysis. Each data point represents a mean and SEM. LI-III = 8 cells, LIV-VI = 6 cells; *** p < 0.001, ** p < 0.01, * p < 0.05

However, it appears that the dendritic spread (Fig. 5F(ii)) for cells in the deep layers is much broader, while for superficial cells, it peaks 100μm from the soma.

### Monosynaptic retrograde labelling of inputs to VIP interneurons in the MEC

To directly assess the structures that send projections to MEC VIP interneurons, we used monosynaptic retrograde rabies virus tracing (Callaway and Luo 2015). To this end, we injected cre-dependent AAV encoding the TVA receptor and rabies glycoprotein into all layers of the MEC in VIP-cre mice, followed by the rabies G virus three weeks later (Fig. 6A). Whole-brain sectioning in coronal and sagittal planes revealed inputs originating within and outside of the MEC. Cells that expressed the TVA-receptor (RFP) and RnvA protein (GFP) were identified as ‘starter’ cells (Fig. 6B(iv & vii)), while those that expressed only the GFP reporter were considered as ‘input’ cells. We excluded one animal as no starter cells were visible. In two out of four animals, we observed input cells within deep layers of the MEC (Fig. 6B(iii)). Comparing our sections with plates from the mouse brain atlas (Paxinos and Franklin, 2001) we found input cells in (i) the mesencephalic trigeminal nucleus (Me5) (Fig. 6C(i), D(ii) and E(i-iii)) and (ii) superior paraolivary complex (SPO) in all four animals (Fig. 6C(ii), D(iii)). Thus, VIP interneurons in the MEC receive local input from within the MEC and long-range input from brainstem (Me5) and midbrain (SPO) nuclei.

**Fig 6.**
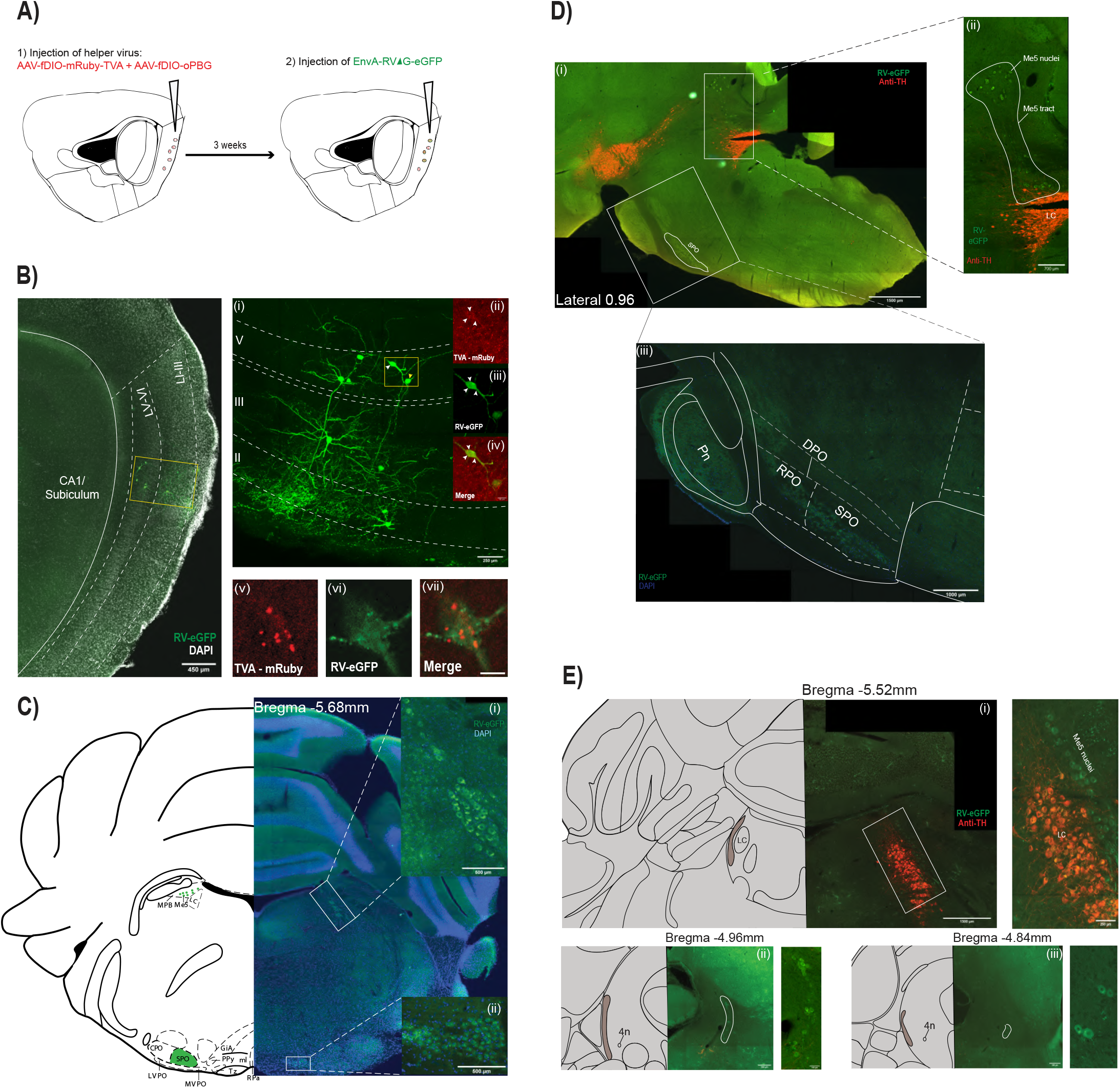
Monosynaptic retrograde tracing in VIP interneurons using rabies viral strategy. (A) Schematic of the tracing strategy. Starter cells or cell bodies infected by the helper and EnvA (RV-eGFP) protein are labelled in red and green respectively. (B) Image of the injection site and starter cells from one animal in 10X magnification. Layers of the MEC are marked by dashed lines.A 20X maginification of ramed area is seen in (i) where the starter and input cells within the MEC are observed and denoted by white and yellow arrows respectively. Transfection of VIP cell by TVA receptors (ii) and (iii) EnvA protein. Coloclaization of the two channels is seen (iv) in the starter cell marked by white arrows where as presenence of TVA is absent in the other cell. 40X Maginifcation of a starter cell as seen from a second animal (v-vii). Presence of colocalization of TVA receptors (v) and EnvA protein (vi) is seen in vii. (C) Monosynaptic inputs to MEC VIP interneurons from the Me5 nuclei (ii) and the SPO (iii) as seen in the coronal plane (D) Sagittal view of the mid-brain and brainstem. (i)Inputs from the Me5 and the SPO are labelled in green. The LC is labelled in red using a stain for tyrosine hydroxylase (TH). 20X magnification of Me5 tract (ii) and overlay of atlas plane to confirm input from (iii)SPO (E) Serial reconstruction of representative sections of Me5 input to. Matching mouse brain atlas planes are aligned with sections. Insets represent a magnified view of the corresponding sections. Me5 – Mesencephalic trigeminal nucleus, SPO – superior paraolivary nucleus, LC – Locus coeruleus.

## Discussion

In this study, we investigate the distribution, morphology, active and passive membrane properties of VIP interneurons in the MEC. We validate that our cre-driver mouse line is efficient and a suitable model system for studying the VIP population in the MEC. Using rabies-mediated viral tracing, we also label the presynaptic partners of VIP MEC cells. Our results confirm that VIPs in the MEC are a heterogenous population of cells, similar to the hippocampus (Tyan et al.,2014), barrel cortex (Prönneke et al. 2015) and the BLA (Rhomberg et al., 2018). Through a combination of electrophysiological, morphological, and viral tracing approaches, this study extends previous findings (Ferrante et al., 2017) in the following ways: We show that there is a gradient in the active and passive membrane properties of VIP cells along the D-V extent of the MEC, and that these differences are preserved between VIP cells in superficial and deep layers. Using retrograde monosynaptic tracing, we reveal that VIP neurons within the MEC receive local inputs from MEC and long-range input from the Me5 and SPO nuclei. These results point to a putative circuit that provides direct information to the MEC about vibrissae movements and auditory localization, respectively.

### VIP cells show heterogeneity in their electrophysiological and morphological properties

Our observations replicate the variability in firing patterns recorded in the hippocampus (Tyan et al., 2014), the barrel cortex (Prönneke et al. 2015) and the BLA (Rhomberg et al., 2018). However, the predominance of IS firing pattern in our dataset differs from some of these previous studies. This increase in IS firing pattern could be explained by the number of CR/VIP cells we observe in the MEC. Previous studies from the hippocampus (Tyan et al., 2014) and the BLA (Rhomberg et al., 2018) report an increase in the incidence of IS firing patterns when the cell coexpresses both VIP and CR. In our data, 67.59% of the total tdTom cells colocalize with CR. Interestingly, we observe that 65% of the total VIP population in the MEC exhibited IS firing patterns. Taking layers into consideration, we see that 79.27% are positive for both CR and VIP markers. This could be reflected by the 76% incidence of IS firing patterns we observe in the superficial layers. Previous studies have noted that VIP cells that colocalize with CR are disinhibitory in nature (Turi et al., 2019; Tyan et al., 2014). While the predominance of CR/VIP in MEC could indicate a primarily disinhibitory circuit motif in the MEC, it remains to be established if our recorded cells with IS firing pattern contained CR. In terms of other electrophysiological properties, our population averages for superficial VIP cells are similar to a prior report, with the exception of a higher percentage of VIP cells with IS firing patterns (Ferrante et al., 2017).

The position of the VIP cell within the MEC is associated with different morphological and electrophysiological properties. In terms of morphological features, we noticed both superficial and deep VIP cells present with similar dendritic and axonal architecture. However, their location in a specific layer dictates their projection patterns. Cells in the superficial layers had either a multi-polar or vertical bi-polar morphology enabling it to span the entire cortical column. These cells are well positioned to receive and transmit incoming information to the entire MEC cortical mantle, while the cells in the deep MEC are found to exclusively cater to the deeper layers of the MEC. Furthermore, we notice that as the cells are located increasingly outwards from the MEC (from LI to LVI), the tendency of the axonal fibers to innervate the superficial layers decreases. Interestingly, this similarity in the neurite processes between superficial and deep MEC is also observed in VIP cells in the barrel cortex (Prönneke et al. 2015). We also identified differences in several electrophysiological properties, notably in input resistance of VIP cell. As a population, MEC VIP cells, are highly excitable, much like their counterparts in other structures (Kepecs & Fishell, 2014; Prönneke et al., 2015; Rhomberg et al., 2018; Tyan et al., 2014). From the current dataset, we found that deep VIP cells are more excitable than superficial cells. Moreover, we saw a gradient in this intrinsic excitability wherein cells located in ventral MEC require less current to fire an action potential compared to cells in dorsal MEC. The gradients in excitability that we report could provide a gating mechanism that prioritizes certain outputs from the MEC as deep MEC sends projections to various intrathalamic structures and LII of the MEC (Ohara et al., 2018; Sürmeli et al., 2015) whereas superficial MEC primarily projects to the hippocampus (Canto et al., 2008). This could also be relevant to the *in vivo* reports that grid cells with smaller spatial scales are present near the dorsal border while cells with a larger spatial scale are found more ventrally (Hafting et al. 2005, Barry et al. 2007, Brun et al. 2008, Stensola et al. 2012).

### VIP cells receive input from within the MEC, the mesencephalic trigeminal nucleus (Me5) and the superior para-olivary complex (SPO)

Our tracing experiments reveal that VIPs receive both local and long-range inputs thus providing for a dynamic routing of information to the MEC. Previous studies have shown that grid cells in LII of the MEC require hippocampal information to stabilize their firing fields (Bonnevie et al., 2013), and that this input is primarily received by cells in the deep layers (Sürmeli et al. 2015, Ohara et al. 2018). While cells from LVb project to the superficial layers (Ohara et al., 2018), our experiments revealed an additional route for this feedback to reach spatially tuned cells in layers II-III. Additionally, VIP interneurons receive inputs from the Me5 an important component of the oromotor neural circuitry, jaw movement and mastication in mice. Afferents from these structures carry information from mechanoreceptors, thermoreceptors and nociceptors that are present in the face and oral cavities (Lazarov, 2007). Additionally, Me5 encodes whisking movements of micro vibrissae (Mameli, Caria, Biagi, Zedda, & Farina, 2017; Mameli et al., 2014), but more specifically the mid-point of whisking (Severson, Xu, Yang, & O’Connor, 2019). Most relevant to the role of the MEC in supporting navigation, whisking mid-point provides the best estimations for localization of objects in space (Cheung, Maire, Kim, Sy, & Hires, 2019), a function that requires stellate cells in LII of the MEC (Tennant et al., 2018). Whisking and movements of oro-facial muscles are behavioural features seen during active exploration and are thus may constitute an important aspect of rodent navigational strategies (Huet and Hartmann 2014; Wills, Muessig, and Cacucci 2014; Lebedev, Pimashkin, and Ossadtchi 2018). Lesions to the Me5 reveal that animals rely on previously acquired navigational strategies and have severe impairment in resetting spatial memory (Ishii et al., 2010). Boundaries of an environment are actively sensed through tactile and whisker movement by mice, and these inputs are thought to be fundamental to place and grid cell firing (Solstad et al., 2008), yet the structure responsible for relaying this information to MEC has largely been unidentified. Our results point to a putative circuit mechanism whereby the Me5 nucleus projects to MEC VIP cells to selectively dis-inhibit MEC border cells when vibrissae contact the walls of the environment.

Along with the Me5, VIP cells also receive dense projections from the superior para-olivary complex (SPO) in the auditory brainstem. The SPO, part of the superior olivary complex, is a GABAergic nuclei (Kulesza, Kadner, & Berrebi, 2007) that receives dense projections form the cochlear nucleus before transmitting this information to the auditory cortex. This structure acts as the focal point for the convergence of binaural signals, where left and right auditory fibers converge (Marrs, Morgan, Howell, Spirou, & Mathers, 2013) thus creating the interaural time difference which aids in localization of sounds by providing a cue to the direction or angle of the source (Beiderbeck et al., 2018). Spatially tuned cells in the MEC can code for non-spatial dimensions such as auditory tone (Aronov, Nevers, & Tank, 2017) and neurons within the MEC respond to loud noise (Zhang et al., 2018). While the source for this auditory input was thought to arise through the medial septum, our rabies tracing experiments reveal a circuit where the MEC receives direct input from the auditory brain stem via potential inhibition of VIP interneurons.

### Functional implications

There is a growing body of work, suggesting that VIP interneurons are critical for facilitating associative learning and maintenance of the E/I balance in various structures. Within the MEC,previous studies have shown the differential inactivation effects of PV or SOM cells on grid and aperiodic cells, respectively. These findings suggest that these interneuron populations may play profoundly different roles in the spatiotemporal control of the local networks within the MEC (Miao et al., 2017). In most structures, VIP cells innervate SOM and PV cells. Our data show that VIP cells in the layers II-III are well-positioned to transmit incoming information to cells across the different layers of the MEC. Considering recent studies, it also appears that grid cells and other spatial cell types in the MEC are modulated by different navigational strategies (Aronov et al., 2017; Butler, Hardcastle, & Giocomo, 2019) and are modulated by sensory inputs, but the source for this input or the circuits required to integrate this information remained unknown (Moser et al., 2014). These findings specify the need for a circuit mechanism in the MEC that would require the consolidation of local and self-motion cues with sensory/modulatory stimuli. Given their connectivity, VIP interneurons can integrate inputs from deep layers and their presynaptic partners in the Me5 and SPO. We predict that these inputs enable VIP cells to provide for a global disinhibitory tone thus creating a temporal window for firing of excitatory cells within the MEC. Our results show that the VIPcre and VIPtdTom mice line are suitable models for future experiments to determine the functions of this cell population *in vivo.*

## Materials and Methods

### Animals

This study was carried out in accordance with the recommendations of the Canadian Council on Animal Care and the McGill University Animal Care Committee. The protocol was approved by the McGill University Animal Care Committee. Animals were housed in a temperature-controlled room with a 12/12 hr light/dark cycle and food and water ad libitum. For this study, we crossed homozygous Vip-ires-cre (VIP^tm1(cre)Zjh^, The Jackson Laboratory, Bar Harbor, ME, USA) mice with homozygous Ai9 lox-stop-lox-tdTomato cre-reporter strain mice (RRID: IMSR JAX:007905) to generate VIPcre/tdTom mice in which tdTom is exclusively expressed in cells that have cre recombinase. tdTom fluorescence was used to identify VIP cells. VIP-ires-cre mice were used for viral tracing experiments.

### Immunohistochemistry

VIPcre/tdTom mice were deeply anesthetized and intracardially perfused with 4% paraformaldehyde in PBS. Brains were dissected, post-fixed by immersion in the same fixative. Free-floating sagittal sections (40 μm) of the entire MEC were cut using a vibratome (VT 1200S, Leica, Wetzlar, Germany). Every fourth ection was collected for IHC. Post slicing, the sections were rinsed in 1X PBS (3X 5 min) and blocked with 1X PBS containing 0.45% fish gelatin and 0.25% Triton-X (3X 15min each). Sections were then incubated with primary antibodies for 48 hr at 4°C on a rotating shaker. The following primary antibodies were used: 1:500 rb-α-VIP Immunostar 722001/20077 (ImmunoStar, Dietzenbach, Germany), 1:500 Ms-α-PV PARV-19 monoclonal P3088 (Millipore Sigma, Saint Louis, Missouri, USA), 1:1000 Ms-α-SOM (H-11) sc-74556 (Santa Cruz Biotechnology, CA, USA), 1:3000 rb-α-Calretinin (CR) 7697 (Swant, Marly, Switzerland) and 1:2000 rb-α-GABA A 2052 (Milipore Sigma). The sections were then subsequently rinsed in 1X PBS (3X 5min) and then incubated in Alexa Flour 488-conjugated AB raised against rabbit IgG and mouse IgG at 1:1000 for 2hrs at RT. After several washes with 0.3% PBS-T the sections were then mounted, and cover slipped with mounting medium Fluoromount-G and DAPI, 0100-20 (SouthernBiotech, Birmingham, AL, USA). For IHC against VIP and CR n = 3 animal and for staining against GABA n = 2 animals. For IHC against PV and SOM n = 1 animals.

#### Rabies circuit mapping

For circuit mapping experiments, brains from injected animals were sliced in the coronal (n = 2) and sagittal plane (n = 3) to retrieve whole brain sections. The following primary antibodies were used to amplify GFP, RFP and stain for tyrosine hydroxylase: 1:1000, rb-α-anti-TH ab112 (Abcam, Cambridge, UK), 1:1000 chk-α-GFP A10262 (ThermoFisher Scientific), 1:1000 rb-α-GFP A11122 (ThermoFisher Scientific), 1:10,000 chk-α-RFP CA600-901-379 (VWR Rockland). Alexa Fluor dyes conjugated to either 488, 555 and 568 were used for secondary antibodies at 1:1000 dilutions.

### Image acquisition and Data analysis for quantification

Quantification of VIP, CR and GABA colocalization was performed using the optical fractionator method in StereoInvestigator software with a Zeiss ApoTome structured illumination device on a widefield microscope. Contours were drawn to delineate the superficial (L1-III) and deep layers (LIV-VI) of the MEC using DAPI as reference. Using live counting, all markers were counted according to optical dissector inclusion-exclusion criteria at each cell’s widest point. Separate markers were used to count the number of tdTom (RFP), VIP/CR/GABA (GFP) and colocalized or double labelled cells (yellow). While LIV is lamina dissecans in the MEC, we repeatedly found many tdTom cell bodies in between LIII (on the outer edge of the layer) and LV as also previously reported in Canto et al., 2008. Hence for all deep cell analysis, VIPtdTom somas found in this region were included in the LIV-LVI category.

### In Vitro Electrophysiology

VIPcre/tdTom (N = 15 animals), of either sex, aged between 5-8 weeks were killed by decapitation and the brain was dissected and placed in an ice cold high-sucrose solution in the following composition: 252 mM sucrose, 24 mM NaHCO3, 10 mM glucose, 3 mM KCl, 2 mM MgCl2, 1.25 mM NaH2PO4, and 1 mM CaCl2, continuously oxygenated with 95% O2/5% CO2;pH 7.3. Sagittal sections of 350 μm were cut according to the method previously described by Pastoll et al (Pastoll, White, & Nolan, 2012). Following this, the sections were transferred to an artificial cerebrospinal fluid (aCSF) solution at RT (solution as above with 126 mM NaCl replacing sucrose, 4.5 mM KCl and 2 mM CaCl2) for 30 min before recording. Slices were recorded in a bath aCSF heated to 35°C. VIPtdTom cells were identified by tdTom fluorescence using an upright BX51WI Olympus microscope with a 60X immersion objective (Olympus, Canada) and an X-cite Series 120Q fluorescence system (Lumen Dynamics). For whole cell patch clamp recordings, glass pipettes of 5-7MΩ resistance were filled with: 144 mM K-gluconate, 10 mM HEPES, 3 mM MgCl2, 2 mM Na2ATP, 0.3 mM GTP, and 0.2 mM EGTA; adjusted to pH 7.2 with KOH. Membrane potentials were not corrected for a junction potential of approximately −10mV. VIPtdTom cells were recorded using a visually guided whole-cell patch clamp technique, a Multiclamp 700B amplifier, a DigiData 1440A digitizer and pClamp10 software.

### Morphology of VIP cells

For the morphology of VIP cells, cells were patched using the above intracellular solution with the addition of neurobiotin (0.4%, SP-1120-50, Vector Laboratories, ON, Canada). To obtain efficient biocytin fills, cells were subjected to a high amount of current at the end of each recording session (Jiang et al., 2015). Following this, the pipette was left undisturbed in whole-cell mode for 30 min to allow for the diffusion of neurobiotin through all neuronal processes. Finally, the pipette was retracted carefully to ensure that the cell membrane did not rupture, as this would result in incomplete fills or the loss of the soma. Slices were fixed in PFA (4% in PBS) at 4°C overnight. Fixed slices were incubated with streptavidin Alexa 488 (1:1000, Jackson ImmunoResearch, West Grove, PA, United States) and NeuroTrace 647 (1:1000) in 0.3% PBS-Triton overnight at room temperature. Slices were mounted on glass slides with Fluoromount G, 0100-01 (Southern Biotech). Biocytin-labeled VIP cells were imaged under the MBF Zeiss ApoTome microscope with a 20X magnification (Carl Zeiss, Germany). The cells were manually reconstructed using Neurolucida 11 software (MBF Bioscience, Williston, United States).

Using cells labelled with NeuroTrace or DAPI as a reference image, contours of the different layers were marked. Dendrites were distinguished from axons by the presence of spines. Initial axon segments were identified by their smooth appearance near the soma which later appeared discontinuous as the axon continued branching. Due to slicing techniques, neuronal processes may be cleaved, leaving abrupt endings visualized as ball-like structures at the end of the processes. The dendrites and axons of filled cells were noted for these abrupt endings and if present were excluded from analysis.

### Monosynaptic circuit mapping using rabies mediated viral tracing

For rabies mediated viral tracing, mice (n = 5 VIP cre) at 5 weeks were anesthetized via inhalation of a combination of oxygen and 5% Isoflurane before being transferred to the stereotaxic frame (David Kopf Instruments), where anesthesia was maintained via inhalation of oxygen and 0.5-2.5% Isoflurane for the duration of the surgery. Body temperature was maintained with a heating pad and eyes were hydrated with gel (Optixcare). Carprofen (10 ml/kg) and saline (0.5 ml) were administered subcutaneously at the beginning of each surgery. All injections were administered via glass pipettes connected to a Nanoject II (Drummond Scientific, PA, USA) injector at a flow rate of 23 nl/sec. For each craniotomy, 2 injection sites from bregma along the medial lateral axis were used, for the first site, the injection pipette was lowered 3.65 mm from the bregma and for the 2^nd^ site it was lowered at 3.45 mm from bregma. For both injection sites 0.260 mm anterior to transverse sinus and an angle of 8° rostral to surface of the brain was used to target the MEC. The depth of the injection site was determined from the extent at which the micropipette bent once it hit the dura lining surface of the brain. Approximate depths for 5-week-old mice were: 1^st^ injection: −2.240, 2^nd^ injection site: −3.00. For each injection site, the pipette was retracted 200 μm after the first bend and the virus was injected at 3 different depth along the dorsal-ventral at 300 μm intervals along extent of the MEC, such that the last injection site usually corresponded to −1.450 to −1.750 along the DV axis. After each injection, the micropipette was left undisturbed for 3-4 min to allow for the infusion of the virus before injecting into the next site. A total of 400 μl of virus was injected in each animal.

#### Injection timeline

Mice were unilaterally injected with a 1:1 combination of AAV.fDIO.mRuby.TVA and AAV.fDIO.oPBG. After allowing for 3 weeks of expression, animals were anesthetized as previously described and EnvA.RV-G.eGFP was injected using scar on skull as a landmark. One-week post injection, animals were sacrificed for quantification of inputs and starter cells. Viral products were a gift from Dr.Byung Kook Lim (University of California San Diego, San Diego, CA, USA)

### Data analysis and statistics

Electrophysiological properties were analyzed with Clampfit10 Software. Recordings were kept for analysis if the spikes overshot 0mV and access resistance was <30MΩ. Membrane resistance (Rm) and access resistance (Ra) were measured in voltage clamp (vc) using pClamp10 software. The following properties were assessed in current clamp (cc). Resting membrane potential (RMP) and spontaneous spiking were assessed over a 30 s recording with no holding current. To assess spike properties, cells were held at a holding potential (h.p.) of −65 mV and a series of 500 ms depolarizing current steps was applied. The step which elicited the first spike was used to assess rheobase, decay tau, spike amplitude and half width, and after-hyperpolarization (AHP) amplitude and time. Firing pattern generated on double the rheobase current were used to categorize the firing patterns based on the Petila convention (Ascoli et al.2008). Spikes elicited for 100pA current injection were used to analyze and determine firing rate, peak interspike interval (ISI) (ISI for the first two spikes), steady state ISI (ISI for the last two spikes), and spike frequency adaptation index[(steady state ISI – peak ISI)/steady state ISI]. To assess the hyperpolarization-activated sag current, if present, a series of hyperpolarizing current steps was applied at a h.p. of −65 mV. The step which hyperpolarized the cell to −120 mV was used to calculate the sag amplitude, measured as the difference between peak and steady state hyperpolarization. Procedures to measure the location of the cell along the dorsal-ventral axis and in a specific layer were modified from (Pastoll et al., 2012). In brief, after each recording session, the pipette would be pulled out of the cell carefully from a whole-cell configuration to a cell-attached mode. Using a low magnification objective (40X) pictures of the position of the pipette relative to the dorsal border of the MEC were taken and stitched together for each cell. This was performed to confirm the location of the recorded cell for a specific layer and its location along the dorsal-ventral axis of the MEC. A total of 51 VIP cells were patched and recorded from.

Properties of 14 (LI-III = 8 cells; LIV-VI = 6 cells) reconstructed neurons were quantified with Neurolucida 11 software (MBF Bioscience, United States). These neurons were considered for reconstruction as they had no visualized swellings in their axonal and dendritic arbours. Dendritic and axonal complexity were assessed using sholl analysis in which concentric sholl segments are generated starting from the center of the cell body (radial interval: 10 μm) and the number of process intersections per sholl segment are recorded. Layer analysis was conducted to estimate the total length of the neural process contained within each layer of the MEC (Neurotrace/DAPI was used to distinguish between the different layers of the MEC). Data were not corrected for tissue shrinkage.

Statistical analysis was carried out using GraphPad prism software (San Diego, CA, USA). Mean values are ± standard error of the mean (SEM). Statistical significance was tested with twotailed Mann-Whitney U test (layer differences, vertical and horizontal extent of neurite process) and liner regression (dorsal-ventral). R^2^ values are stated for results of liner regression. A two-way analysis of variance (ANOVA) was performed to determine the length of neurite process present within in a layer. A *post hoc* Bonferroni test on all pairwise comparisons was performed to determine significant different bins.

## Acknowledgments

We thank Drs. Coralie-anne Mosser, Jennifer C. Robinson, Alexandra T. Keinath and Robert R. Rozeske for comments on earlier versions of this manuscript. The present study used the services of the Molecular and Cellular Microscopy Platform in the Douglas Hospital Research Center. This work was supported by a McGill University ‘Healthy Brain for Healthy Lives’ CFREF doctoral fellowship to S.B., and funding from the Canadian Institutes for Health Research grants (#367017 and #377074), the National Sciences and Engineering Council (Discovery grant #74105), the Scottish Rite Charitable Foundation, and the Canada Research Chairs Program to M.P.B.

## Author contributions

All authors helped to conceive and plan this study. S.B. collected and analyzed all data. B.K. provided help to plan and execute the rabies tracing experiments. S.B. and M.P.B wrote the manuscript.

## Competing interests

None.

---

**Figure supplement 1 (S1).**
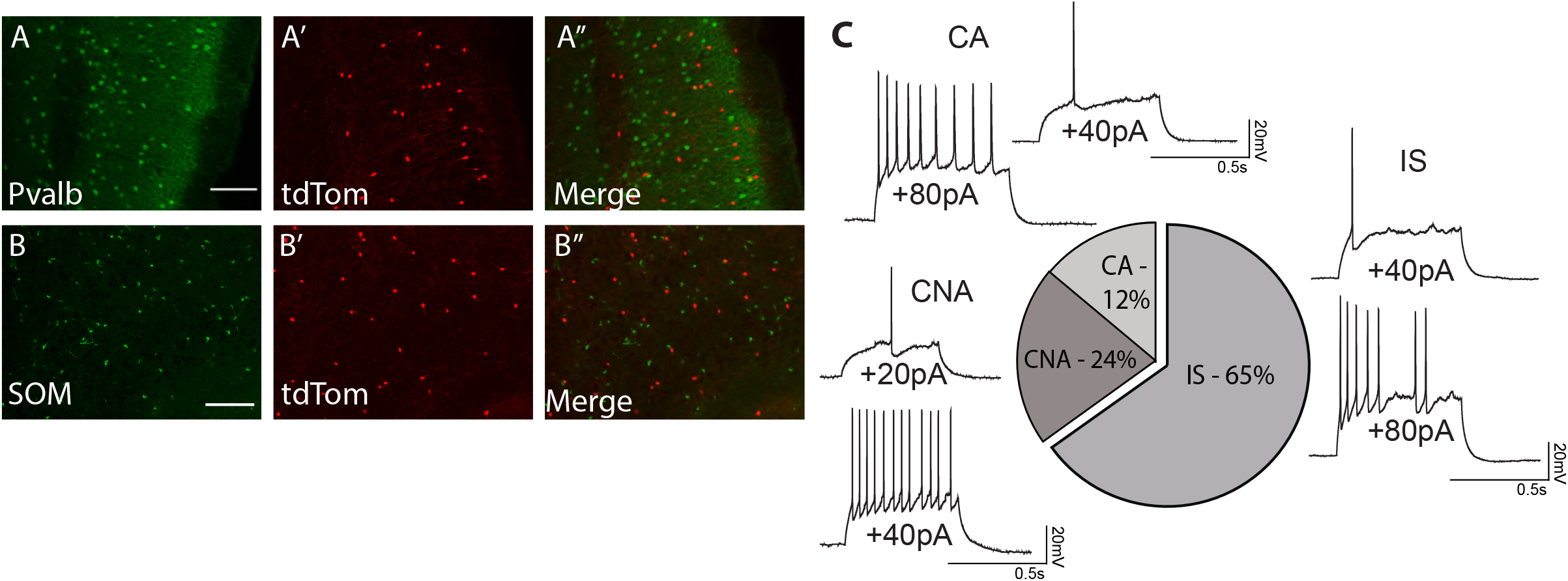
(A-B”) Maximum projection intensity of a 40-μm thick sagittal section of superficial MEC treated with IHC using αPV AB (A-JA”) and αSst AB (B-B”). Scale bar is 250μm (C) Percentage of action potential firing patterns observed in VIP cells in the MEC. Examples of membrane potential responses to current steps recorded at rheobase and suprathreshold current. Abbreviations name the firing pattern after the Petila convention (Ascoli et al.2008; IS, Irregular spiking; CA, Continuous adapting; CNA, continuous non-adapting).

## Notes

### Competing Interest Statement

The authors have declared no competing interest.

